# Computational modeling of molecular structure of microRNA inhibitors selected against microRNA over-expressed in thyroid cancer

**DOI:** 10.1101/2021.04.24.441267

**Authors:** Luís Jesuíno de Oliveira Andrade, Alcina Maria Vinhaes Bittencourt, Luís Matos de Oliveira, Gabriela Correia Matos de Oliveira

## Abstract

**Introduction:** Thyroid cancer is the most prevalent malignant neoplasm of endocrine system and advances in thyroid molecular biology studies demonstrate that microRNAs (miRNAs) seem to play a fundamental role in tumor triggering and progression. The miRNAs inhibitors are nucleic acid-based molecules that blockade miRNAs function, making unavailable for develop their usual function, also acting as gene expression controlling molecules.

**Objective:** To develop *in silico* projection of molecular structure of miRNA inhibitors against miRNA over-expressed in thyroid cancer.

**Methods:** We conducted a search of the nucleotide sequence of 12 miRNAs already defined as inhibitors against miRNA over-expressed in thyroid cancer, realizing *in silico* projection of the molecular structure of following miRNAs: miRNA-101, miRNA-126, miRNA-126-3p, miRNA-141, miRNA-145, miRNA-146b, miRNA-206, miRNA-3666, miRNA-497, miRNA-539, miRNA-613, and miRNA-618. The nucleotides were selected using GenBank that is the NIH genetic sequence database. The sequences obtained were aligned with the Clustal W multiple alignment algorithms. For the molecular modeling, the structures were generated with the RNAstructure, a fully automated miRNAs structure modelling server, accessible via the Web Servers for RNA Secondary Structure Prediction.

**Results:** We demonstrated a search for nucleotide sequence and the projection of the molecular structure of the following miRNA inhibitors against miRNA over-expressed in thyroid cancer: miRNA-101, miRNA-126, miRNA-126-3p, miRNA-141, miRNA-145, miRNA-146b, miRNA-206, miRNA-3666, miRNA-497, miRNA-539, miRNA-613, and miRNA-618.

**Conclusion:** In this study we show *in silico* secondary structures projection of selected of 12 miRNA inhibitors against miRNA over-expressed in thyroid cancer through computational biology.

## INTRODUCTION

Thyroid cancer is the most prevalent malignant neoplasm of endocrine system, originating from cells of different origin of own thyroid gland and although its etiology isn’t yet completely understood it has been observed that many genetic factors would be also involved in their onset.^(1)^ The microRNAs (miRNAs) seem to play fundamental role in tumor triggering and progression of multiples types of cancers, functioning as tumor suppressor genes or oncogenes in several tumors. Most recently, studies has been suggested the involvement of miRNAs as diagnostic indicators of thyroid cancer.^(2)^

The miRNAs have between 19 and 25 nucleotides and are small noncoding RNAs that implicate in post transcriptional control of gene expression in multicellular organisms by disturbing the stability in processes such as translation, having as result target miRNAs degradation or silencing.^(3)^

The miRNA inhibitors are nucleic acid-based molecules that blockade miRNA role. Synthetic miRNA inhibitors integrate the reverse complement of mature miRNA, linking to endogenous matured miRNAs, making it unavailable for develop their usual function.^(4)^

In just over ten years, our knowledge of the structure and role of miRNA has significantly increased. The bioinformatics programs currently available to construct molecular modeling and analyses nucleotide sequences provide tools for assembly of miRNA inhibitors of the thyroid cancer and understanding of their molecular mechanisms. The aim of this study was to develop *in silico* projection of molecular structure of 12 miRNAs already defined as inhibitors against miRNA over-expressed in thyroid cancer.

## METHODS

We conducted a search of the nucleotide sequence of miRNAs already defined as inhibitors against miRNA over-expressed in thyroid cancer, realizing *in silico* projection of the molecular structure of following miRNAs: miRNA-101, miRNA-126, miRNA-126-3p, miRNA-141, miRNA-145, miRNA-146b, miRNA-206, miRNA-3666, miRNA-497, miRNA-539, miRNA-613, and miRNA-618.

The nucleotide sequences were selected using GenBank that is the NIH genetic sequence database. GenBank is a component of the International Nucleotide Sequence Database Collaboration that includes the GenBank at NCBI, the European Nucleotide Archive, and the DNA DataBank of Japan. The sequences obtained were aligned with the Clustal W multiple alignment algorithms. For the molecular modeling, the structures were generated with the RNAstructure that is a fully automated miRNA structural modelling server, accessible via Web Servers for RNA Secondary Structure Prediction (http://rna.urmc.rochester.edu/RNAstructureWeb/).

### Nucleotide database search and sequence analysis

GenBank is a nucleotide sequence analysis tool available in the public domain (https://www.ncbi.nlm.nih.gov/genbank/submit/) and a wide variety of nucleotide algorithms is used to search many different sequence databases. The Nucleotide, Genome Survey Sequence (GSS), and Expressed Sequence Tag (EST) database all contains nucleic acid sequences. The information in EST and GSS is from two major mass-sequencing distributions of GenBank.

### Building molecular models

The structure and function of amino acid and proteins are determined by their nucleotide sequences and the structure prediction still remains a significant challenge, with a great demand for high resolution structure prediction methods.

Homology modeling is at the moment the higher precision computational method to create trustworthy structural models and is usually utilized in numerous biological applications.

### Modeling with RNAstructure

The Predict to Secondary Structure server calculate a partition function, predict the maximum free energy structure, find structures with maximum expected accuracy, and pseudoknot prediction. This server produces a group of secondary structures with strongly possible similarities annotations, starting with the lowest free energy structure and including others with varying probabilities of correctness. Various structures are incorporated because the minimum free energy structure may not be appropriate. If shape constraints are specified, the shape constraints are applied to the probability annotated structures. In addition, a second group of shape constrained, shape annotated structures will be generated. This shape structure group is distinct from the probability annotated structure group, and is not likely annotated itself.

## RESULTS

We demonstrated a search for nucleotide sequence and the projection of the molecular structure of the following miRNA inhibitors against miRNA over-expressed in thyroid cancer: miRNA-101, miRNA-126, miRNA-126-3p, miRNA-141, miRNA-145, miRNA-146b, miRNA-206, miRNA-3666, miRNA-497, miRNA-539, miRNA-613, and miRNA-618.

### Nucleotide sequence of miRNA-101

The main data source used for reconstructing the miRNA-101 was the nucleotide sequence file in FASTA format. The full-length nucleotide of miRNA-101 was obtained from the GenBank database under the identifier NCBI: NR_029516.1. The miRNA-101 was predicted to encode a 75 bp linear. All coding sequences were selected and exported as nucleotides in FASTA format, using the annotation of the NCBI - Graphics. Homo sapiens miRNA-101 (MIR101) analysis is shown in **Figure 1**.

**Figure 1.**
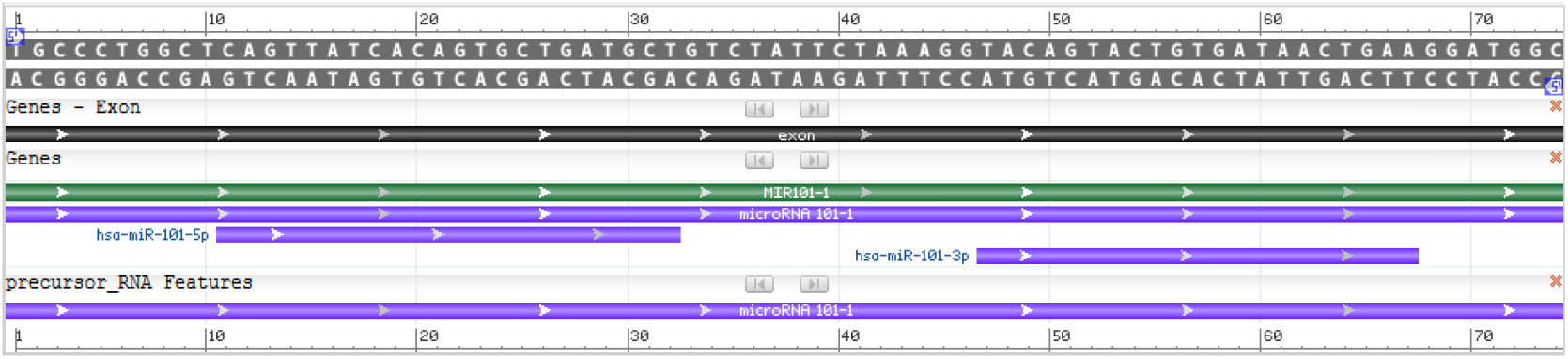
Homo sapiens miRNA-101 (MIR101) - *model-template aligment*. Source: http://www.ncbi.nlm.nih.gov/genbank/

### Molecular model of miRNA-101

Nucleotide sequences of Homo sapiens miRNA-101 (MIR101) were obtain using FASTA format; modeling was conducted using the RNAstructure programs, which were adjusted and optimized for alignment between miRNA-101 nucleotide and structural templates. On the basis of a sequence alignment between the miRNA-101 nucleotide and the template structure, a structural model for the target nucleotide was generated. Appraisal tools were utilized to calculate the trustworthiness of the obtained model. Thus, using the RNAstructure programs automated comparative nucleotide modeling server, we constructed a homology model of the Homo sapiens miRNA-101 (MIR101) (**Figure 2**).

**Figure 2.**
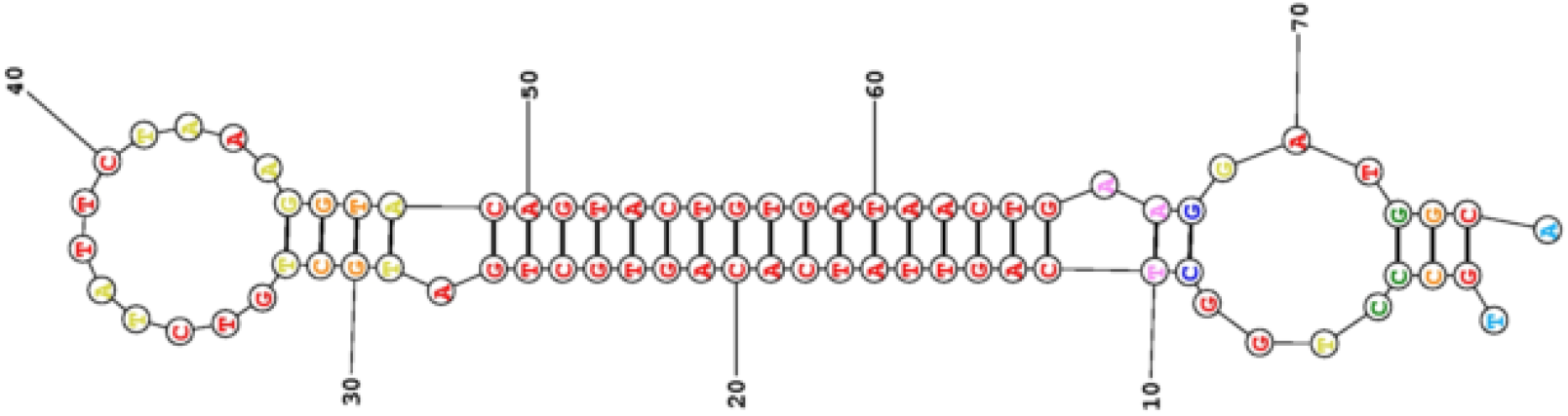
Homology model of the Homo sapiens miRNA-101. Source: Search result - http://rna.urmc.rochester.edu/RNAstructureWeb/

### Nucleotide sequence of miRNA-126

The reconstruction of the miRNA-126 was made from a nucleotide sequence file in FASTA format. The full-length nucleotide of miRNA-126 was obtained from the GenBank database under the identifier NCBI Reference Sequence: NR_029695.1. The miRNA-126 was predicted to encode a 85 bp linear. All coding sequences were selected and exported as nucleotides in FASTA format, using the annotation of the NCBI - Graphics. Homo sapiens miRNA-126 (MIR126), MiRNA analysis is shown in **Figure 3**.

**Figure 3.**
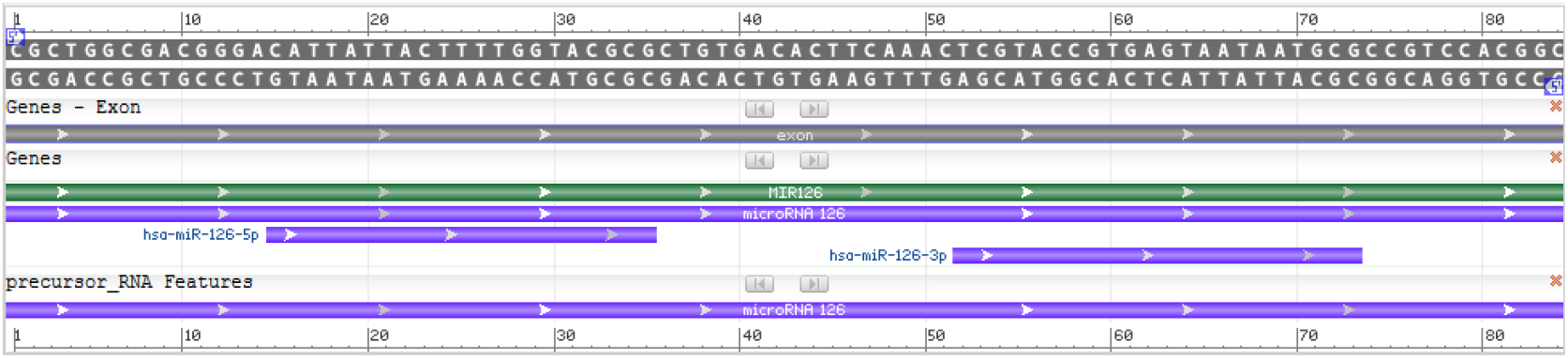
Homo sapiens miRNA-126 (MIR126) - – *model-template aligment*. Source: http://www.ncbi.nlm.nih.gov/genbank/

### Molecular model of miRNA-126

Template search with FASTA format was performed against the RNAstructure template library, which were adjusted and optimized for alignment between Homo sapiens miRNA-126 (MIR126) nucleotide and structural templates. On the basis of a sequence alignment between the miRNA-126 nucleotide and the template structure, a structural model for the target nucleotide was generated. Appraisal tools were utilized to calculate the trustworthiness of the obtained model. According to the described criteria, a model for the Homo sapiens miRNA-126 (MIR126) was generated (**Figure 4**).

**Figure 4.**
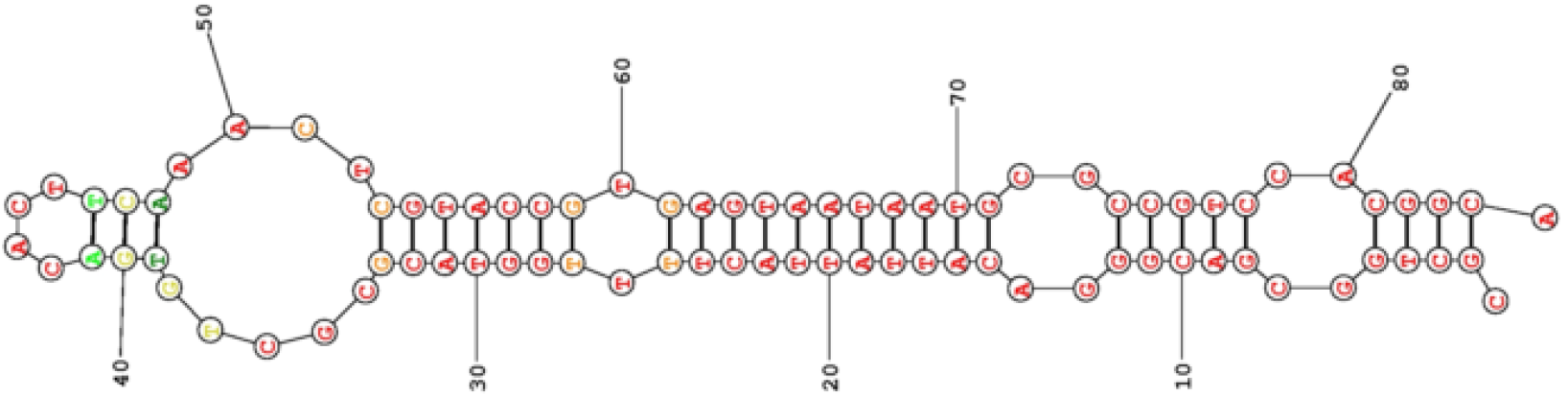
Homology model of the Homo sapiens miRNA-126 (MIR126). Source: Search result - http://rna.urmc.rochester.edu/RNAstructureWeb/

### Nucleotide sequence of miRNA-126-3p

The main data source used for reconstructing the miRNA-126-3p was the nucleotide sequence file in FASTA format. The full-length nucleotide of miRNA-126-3p was obtained from the GenBank database under the identifier Homo sapiens miRNA hsa-miR-126-3p GenBank: LM379063.1. The miRNA-126-3p was predicted to encode a 22 bp linear transcribed-RNA. All coding sequences were selected and exported as nucleotides in FASTA format, using the annotation of the NCBI - Graphics. Homo sapiens miRNA hsa-miR-126-3p analysis is shown in **Figure 5**.

**Figure 5.**
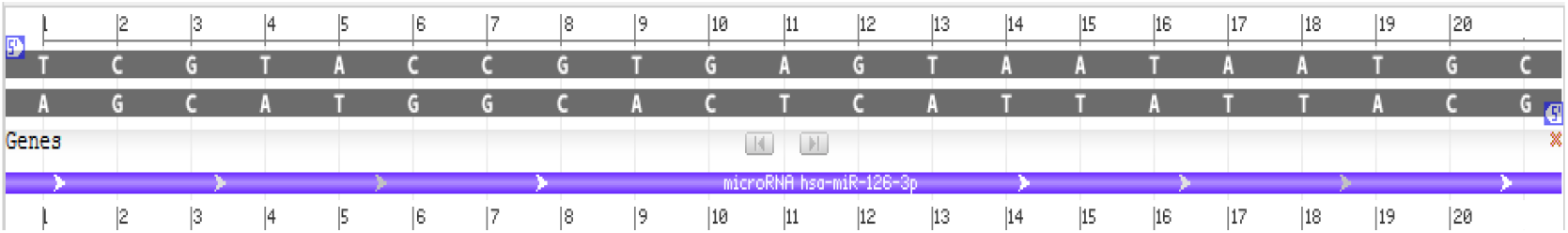
Homo sapiens miRNA hsa-miR-126-3p – *model-template aligment*. Source: http://www.ncbi.nlm.nih.gov/genbank/

### Molecular model of miRNA-126-3p

Nucleotide sequences of miRNA-126-3p were obtain using FASTA format; modeling was conducted using the RNAstructure programs, which were adjusted and optimized for alignment between Homo sapiens miRNA hsa-miR-126-3pnucleotide and structural templates. On the basis of a sequence alignment between the Homo sapiens miRNA hsa-miR-126-3pnucleotide and the template structure, a structural model for the target nucleotide was generated. Appraisal tools were utilized to calculate the trustworthiness of the obtained model. Thus, using the RNAstructure programs automated comparative nucleotide modeling server, we constructed a homology model of the Homo sapiens miRNA hsa-miR-126-3p (**Figure 6**).

**Figure 6.**
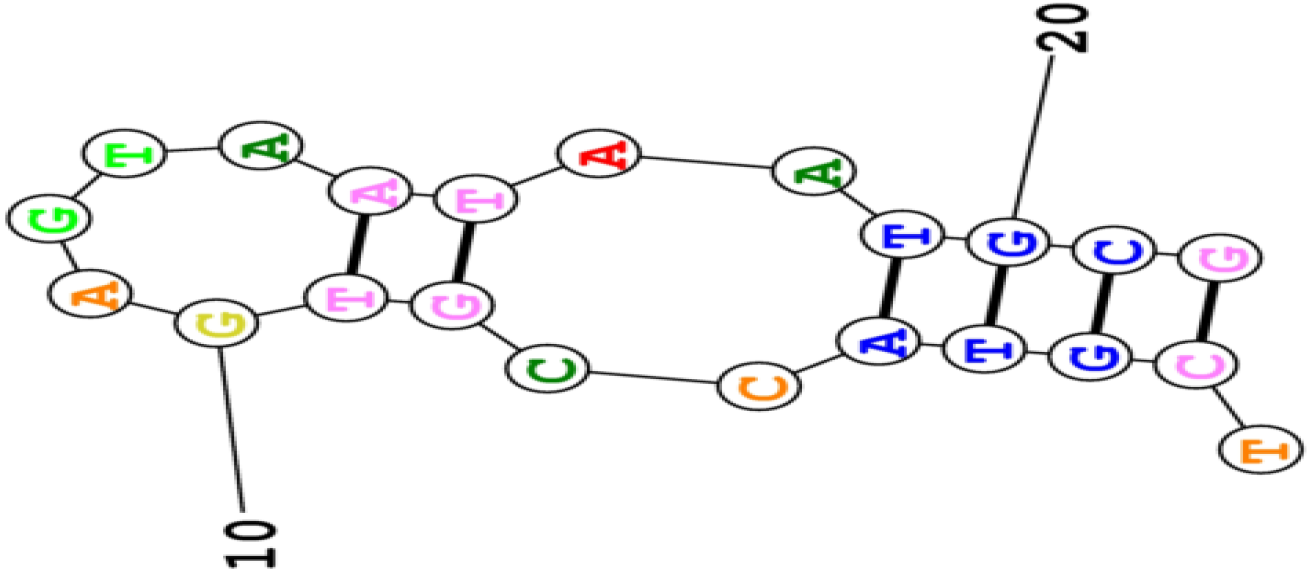
Homology model of the Homo sapiens miRNA hsa-miR-126-3p Source: Search result - http://rna.urmc.rochester.edu/RNAstructureWeb/

### Nucleotide sequence of miRNA-141

The reconstruction of the miRNA-141 was made from a nucleotide sequence file in FASTA format. The full-length nucleotide of miRNA-141 was obtained from the GenBank database under the identifier NCBI Reference Sequence: NR_029682.1. The Homo sapiens miRNA-141 (MIR141), miRNA was predicted to encode a 95 bp linear non-coding RNA, miRNA. All coding sequences were selected and exported as nucleotides in FASTA format, using the annotation of the NCBI - Graphics. Homo sapiens miRNA-141 (MIR141) analysis is shown in **Figure 7**.

**Figure 7.**
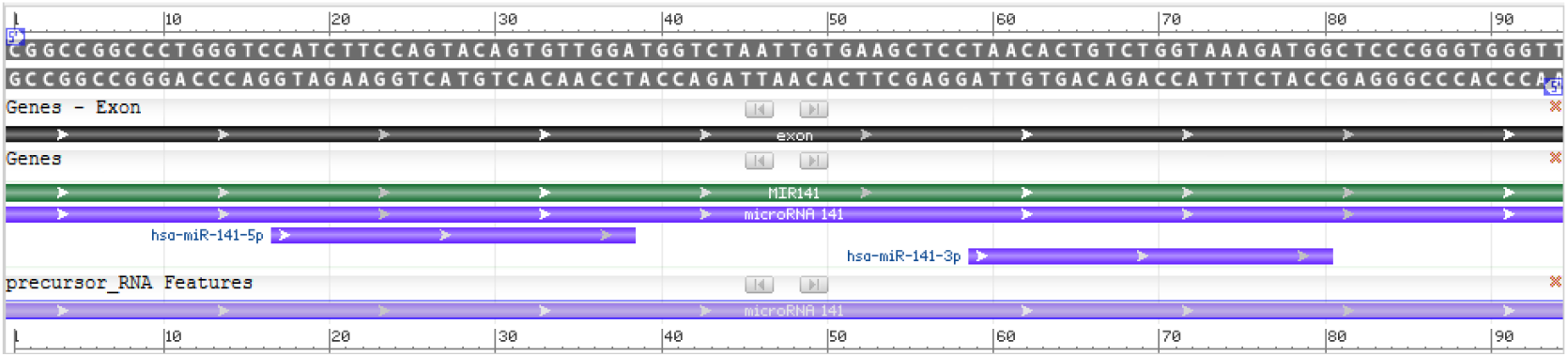
Homo sapiens miRNA-141 (MIR141) - *model-template aligment*. Source: http://www.ncbi.nlm.nih.gov/genbank/

### Molecular model of miRNA-141

Template search with FASTA format was performed against the RNAstructure template library, which were adjusted and optimized for alignment between Homo sapiens miRNA-141 (MIR141), miRNA nucleotide and structural templates. On the basis of a sequence alignment between the miRNA-141 nucleotide and the template structure, a structural model for the target nucleotide was generated. Appraisal tools were utilized to calculate the trustworthiness of the obtained model. According to the described criteria, a model for the Homo sapiens miRNA-141(MIR141), miRNA was generated (**Figure 8**).

**Figure 8.**
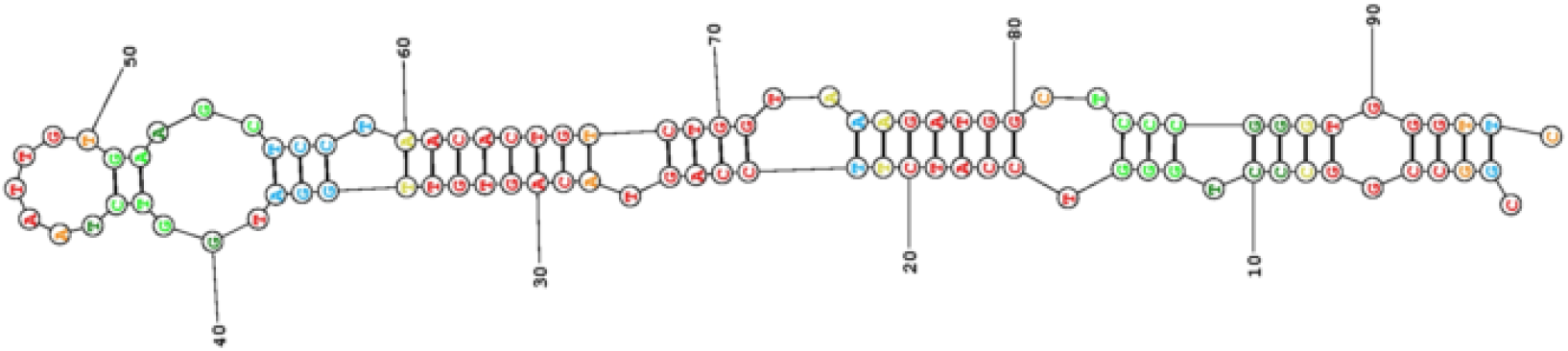
Homology model of the Homo sapiens miRNA-141 (MIR141). Source: Search result - http://rna.urmc.rochester.edu/RNAstructureWeb/

### Nucleotide sequence of miRNA-145

The reconstruction of the miRNA-145 was made from a nucleotide sequence file in FASTA format. The full-length nucleotide of miRNA-145 was obtained from the GenBank database under the identifier NCBI Reference Sequence: NR_029686.1. The Homo sapiens miRNA-145 (MIR145), miRNA was predicted to encode a 88 bp linear non-coding RNA, miRNA. All coding sequences were selected and exported as nucleotides in FASTA format, using the annotation of the NCBI - Graphics. Homo sapiens miRNA-145 (MIR145), miRNA analysis is shown in **Figure 9**.

**Figure 9.**
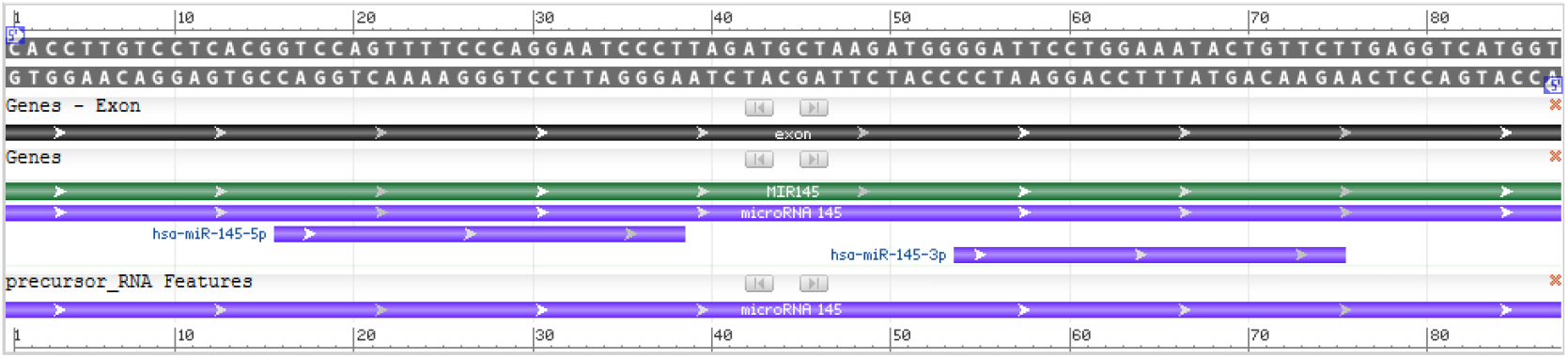
Homo sapiens miRNA-145 (MIR145) - *model-template aligment*. Source: http://www.ncbi.nlm.nih.gov/genbank/

### Molecular model of miRNA-145

Nucleotide sequences of mirRNA-145 were obtain using FASTA format; modeling was conducted using the RNAstructure programs, which were adjusted and optimized for alignment between Homo sapiens miRNA-145 (MIR145), miRNA nucleotide and structural templates. On the basis of a sequence alignment between the Homo sapiens miRNA-145 (MIR145), miRNA nucleotide and the template structure, a structural model for the target nucleotide was generated. Appraisal tools were utilized to calculate the trustworthiness of the obtained model. Thus, using the RNAstructure programs automated comparative nucleotide modeling server, we constructed a homology model of the Homo sapiens miRNA-145 (MIR145), miRNA (**Figure 10**).

**Figure 10.**
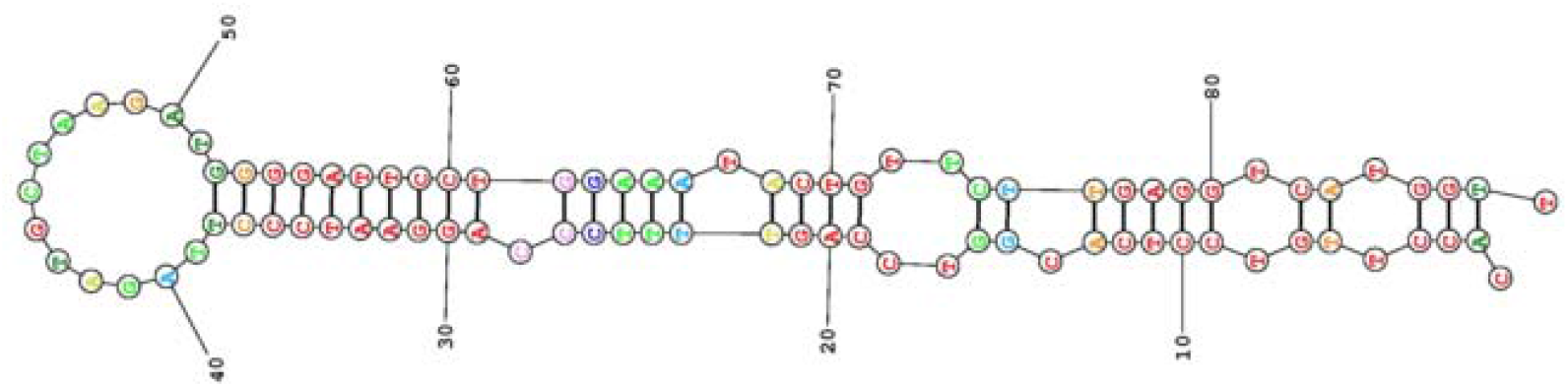
Homology model of the Homo sapiens miRNA-145 (MIR145). Source: Search result - http://rna.urmc.rochester.edu/RNAstructureWeb/

### Nucleotide sequence of miRNA-146b

The main data source used for reconstructing the miRNA-146b was the nucleotide sequence file in FASTA format. The full-length nucleotide of miRNA-146b was obtained from the GenBank database under the identifier Homo sapiens miRNA-146b (MIR146B), miRNA GenBank: NR_030169.1. The miRNA-146b was predicted to encode a 73 bp linear non-coding RNA, miRNA. All coding sequences were selected and exported as nucleotides in FASTA format, using the annotation of the NCBI - Graphics. Homo sapiens miRNA-146b (MIR146B) analysis is shown in **Figure 11**.

**Figure 11.**
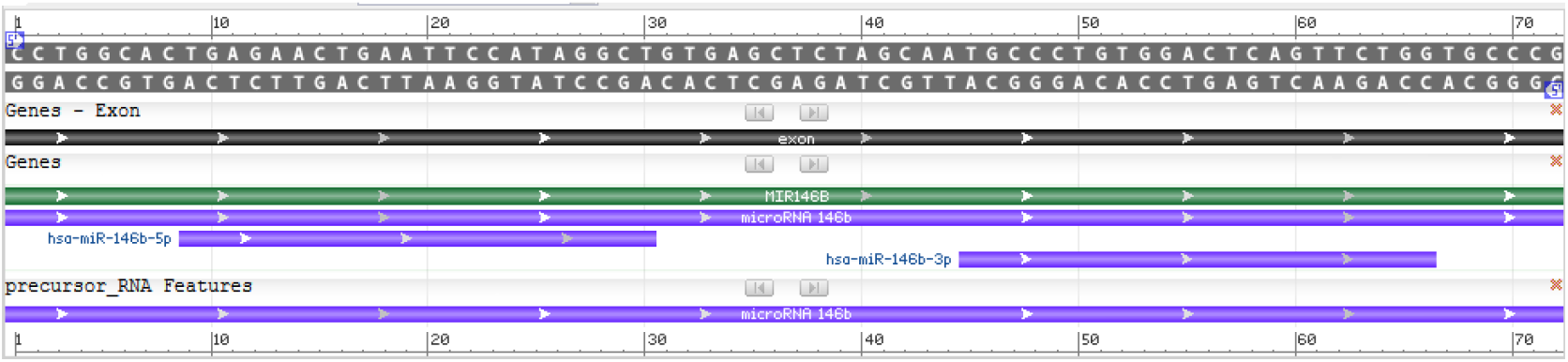
Homo sapiens miRNA-146b (MIR146B) - *model-template aligment*. Source: http://www.ncbi.nlm.nih.gov/genbank/

### Molecular model of miRNA-146b

Template search with FASTA format was performed against the RNAstructure template library, which were adjusted and optimized for alignment between Homo sapiens miRNA-146b (MIR146B), miRNA, nucleotide and structural templates. On the basis of a sequence alignment between the miRNA-141 nucleotide and the template structure, a structural model for the target nucleotide was generated. Appraisal tools were utilized to calculate the trustworthiness of the obtained model. According to the described criteria, a model for the Homo sapiens miRNA-146b (MIR146B), miRNA was generated (**Figure 12**).

**Figure 12.**
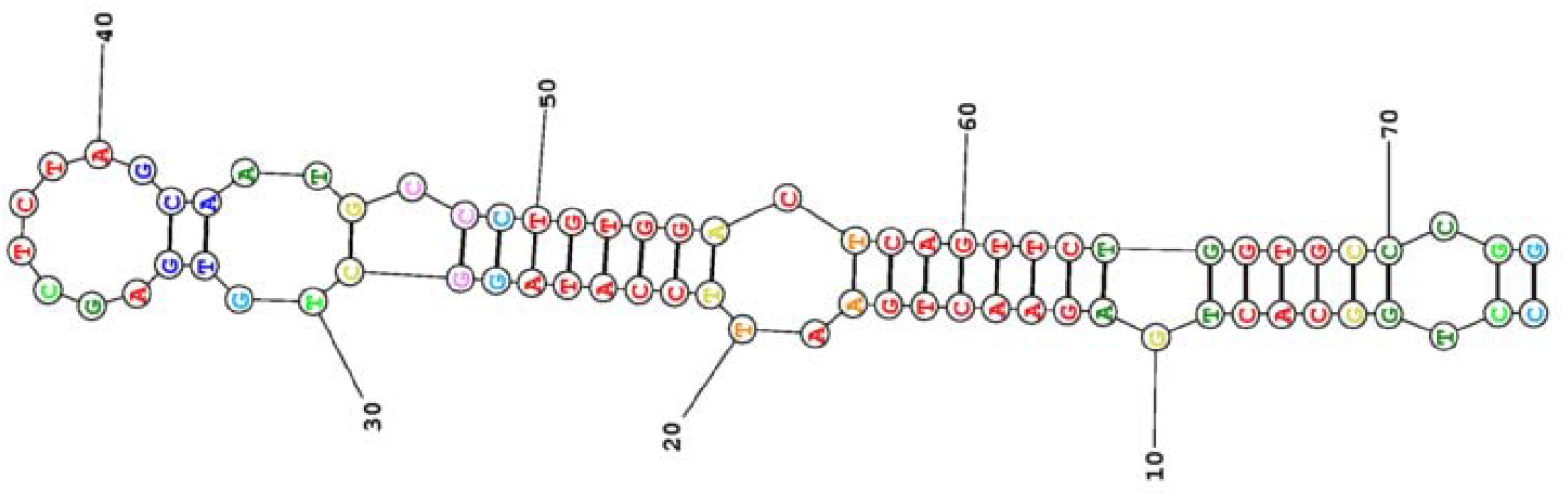
Homology model of the Homo sapiens miRNA-146b (MIR146B). Source: Search result - http://rna.urmc.rochester.edu/RNAstructureWeb/

### Nucleotide sequence of miRNA-206

The main data source used for reconstructing the miRNA-206 was the nucleotide sequence file in FASTA format. The full-length nucleotide of miRNA-206 was obtained from the GenBank database under the identifier NCBI: NR_029713.1. The miRNA-206 was predicted to encode a 86 bp linear non-coding RNA, miRNA. All coding sequences were selected and exported as nucleotides in FASTA format, using the annotation of the NCBI - Graphics. Homo sapiens miRNA-206 (MIR206), miRNA analysis is shown in **Figure 13**.

**Figure 13.**
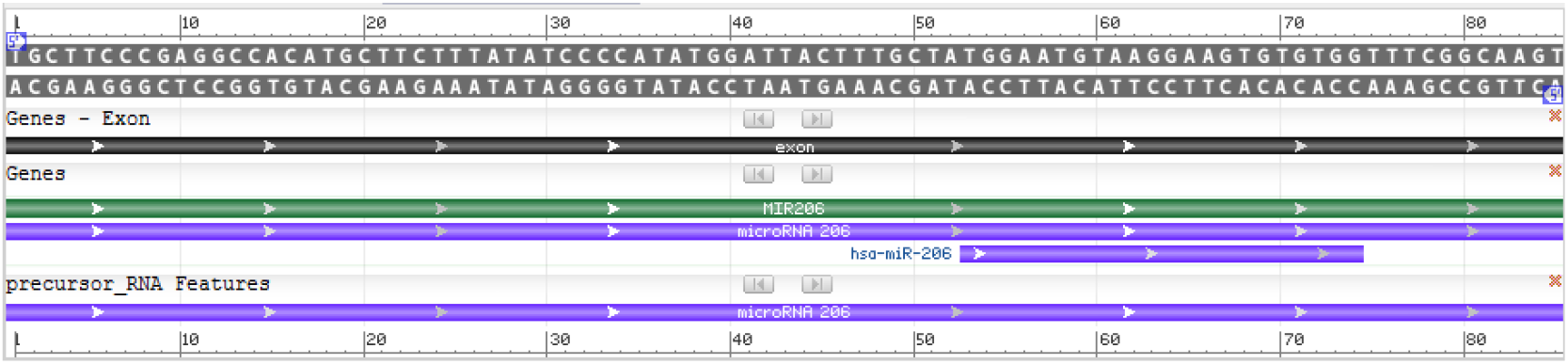
Homo sapiens miRNA-206 (MIR206) - *model-template aligment*. Source: http://www.ncbi.nlm.nih.gov/genbank/

### Molecular model of miRNA-206

Template search with FASTA format was performed against the RNAstructure template library, which were adjusted and optimized for alignment between Homo sapiens miRNA-206 (MIR206), miRNA, nucleotide and structural templates. On the basis of a sequence alignment between the miRNA-206 nucleotide and the template structure, a structural model for the target nucleotide was generated. Appraisal tools were utilized to calculate the trustworthiness of the obtained model. According to the described criteria, a model for the Homo sapiens miRNA-206 (MIR206), miRNA was generated (**Figure 14**).

**Figure 14.**
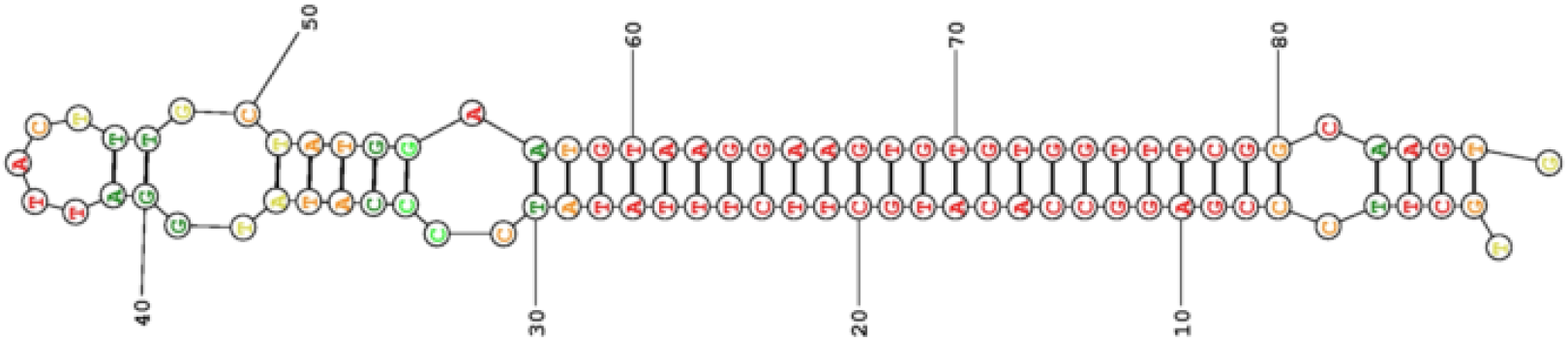
Homology model of the Homo sapiens miRNA-206 (MIR206). Source: Search result - http://rna.urmc.rochester.edu/RNAstructureWeb/

### Nucleotide sequence of miRNA-3666

The reconstruction of the miRNA-3666 was made from a nucleotide sequence file in FASTA format. The full-length nucleotide of miRNA-3666 was obtained from the GenBank database under the identifier NCBI Reference Sequence: NR_037439.1. The Homo sapiens miRNA-3666 (MIR3666), miRNA was predicted to encode a 88 bp linear non-coding RNA, miRNA. All coding sequences were selected and exported as nucleotides in FASTA format, using the annotation of the NCBI - Graphics. Homo sapiens miRNA-3666 (MIR3666), miRNA analysis is shown in **Figure 15**.

**Figure 15.**
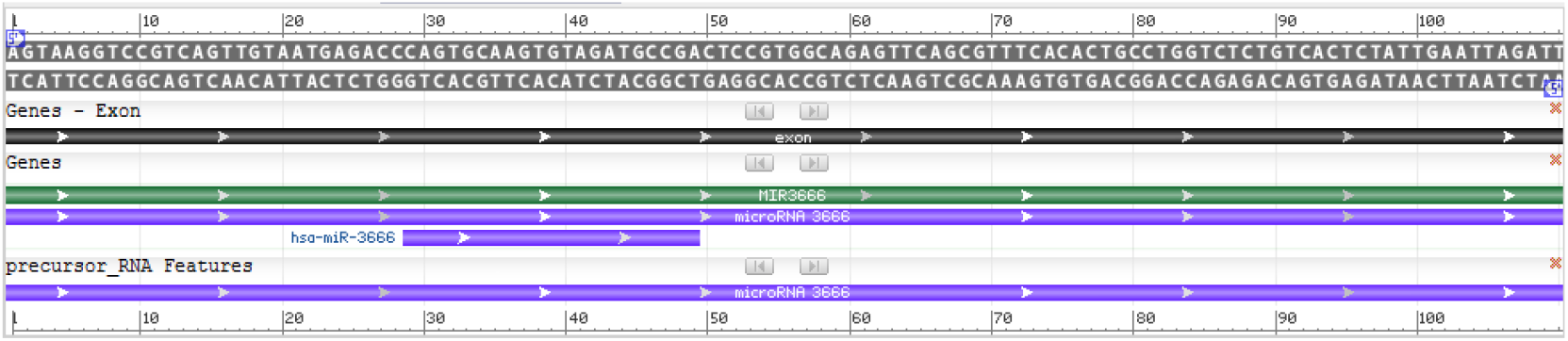
Homo sapiens miRNA-3666 (MIR3666) - *model-template aligment*. Source: http://www.ncbi.nlm.nih.gov/genbank/

### Building a molecular model of miRNA-3666

Nucleotide sequences of miRNA-3666 were obtain using FASTA format; modeling was conducted using the RNAstructure programs, which were adjusted and optimized for alignment between Homo sapiens miRNA-3666 (MIR3666), miRNA nucleotide and structural templates. On the basis of a sequence alignment between the Homo sapiens miRNA-3666 (MIR3666), miRNA nucleotide and the template structure, a structural model for the target nucleotide was generated. Appraisal tools were utilized to calculate the trustworthiness of the obtained model. Thus, using the RNAstructure programs automated comparative nucleotide modeling server, we constructed a homology model of the Homo sapiens miRNA-3666 (MIR3666), miRNA (**Figure 16**).

**Figure 16.**
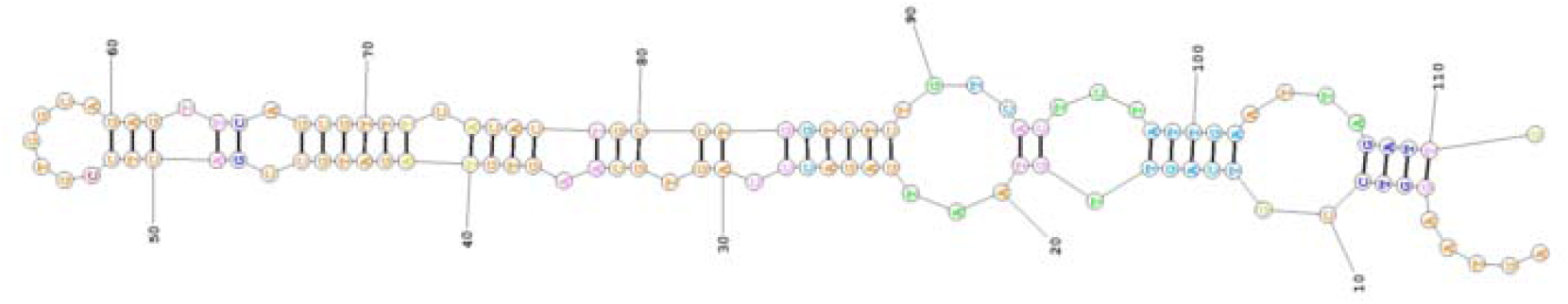
Homology model of the Homo sapiens miRNA-3666 (MIR3666). Source: Search result - http://rna.urmc.rochester.edu/RNAstructureWeb/

### Search for nucleotide sequence of miRNA-497

The main data source used for reconstructing the miRNA-497 was the nucleotide sequence file in FASTA format. The full-length nucleotide of miRNA-497 was obtained from the GenBank database under the identifier Homo sapiens miRNA-497 (MIR497), miRNA GenBank: NR_030178.1. The miRNA-497 was predicted to encode a 112 bp linear non-coding RNA, miRNA. All coding sequences were selected and exported as amino acids in FASTA format, using the annotation of the NCBI - Graphics. Homo sapiens miRNA-497 (MIR497), miRNA analysis is shown in **Figure 17**.

**Figure 17.**
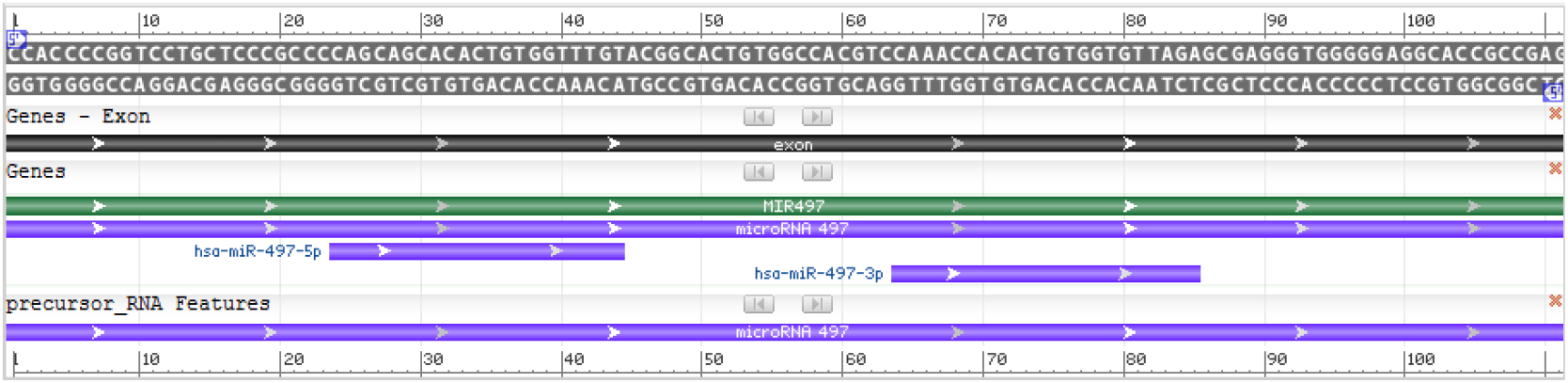
Homo sapiens miRNA-497 (MIR497) - *model-template aligment*. Source: http://www.ncbi.nlm.nih.gov/genbank/

### Building a molecular model of miRNA-497

Template search with FASTA format was performed against the RNAstructure template library, which were adjusted and optimized for alignment between Homo sapiens miRNA-497 (MIR497), miRNA, nucleotide and structural templates. On the basis of a sequence alignment between the miRNA-497 nucleotide and the template structure, a structural model for the target nucleotide was generated. Appraisal tools were utilized to calculate the trustworthiness of the obtained model. According to the described criteria, a model for the Homo sapiens miRNA-497 (MIR497), miRNA was generated (**Figure 18**).

**Figure 18.**
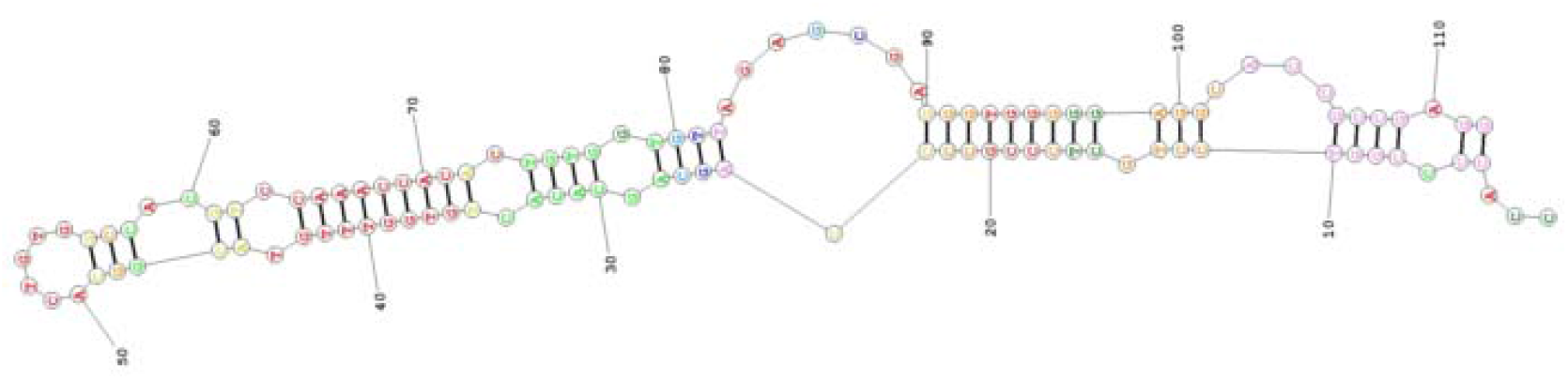
Homology model of the Homo sapiens miRNA-497 (MIR497). Source: Search result - http://rna.urmc.rochester.edu/RNAstructureWeb/

### Search for nucleotide sequence of miRNA-539

The reconstruction of the miRNA-539 was made from a nucleotide sequence file in FASTA format. The full-length nucleotide of miRNA-539 was obtained from the GenBank database under the identifier NCBI Reference Sequence: NR_030256.1. The Homo sapiens miRNA-539 (MIR539), miRNA was predicted to encode a 78 bp linear non-coding RNA, miRNA. All coding sequences were selected and exported as amino acids in FASTA format, using the annotation of the NCBI - Graphics. Homo sapiens miRNA-539 (MIR539), miRNA analysis is shown in **Figure 19**.

**Figure 19.**
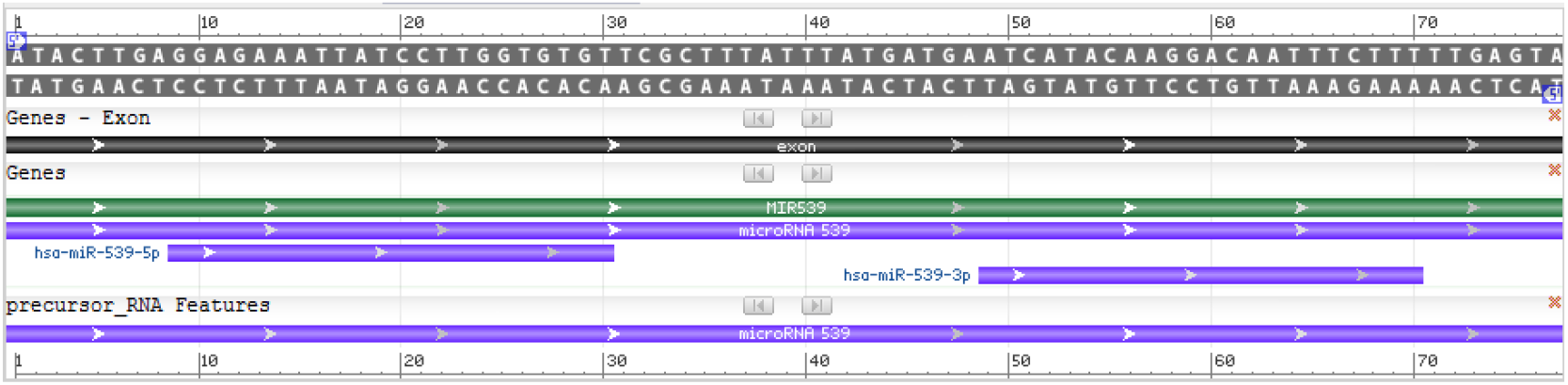
Homo sapiens miRNA-539 (MIR539) - *model-template aligment*. Source: http://www.ncbi.nlm.nih.gov/genbank/

### Building a molecular model of miRNA-539

Nucleotide sequences of miRNA-539 were obtain using FASTA format; modeling was conducted using the RNAstructure programs, which were adjusted and optimized for alignment between Homo sapiens miRNA-539 (MIR539), miRNA nucleotide and structural templates. On the basis of a sequence alignment between the Homo sapiens miRNA-539 (MIR539), miRNA nucleotide and the template structure, a structural model for the target nucleotide was generated. Appraisal tools were utilized to calculate the trustworthiness of the obtained model. Thus, using the RNAstructure programs automated comparative nucleotide modeling server, we constructed a homology model of the Homo sapiens miRNA-539 (MIR539), miRNA (**Figure 20**).

**Figure 20.**
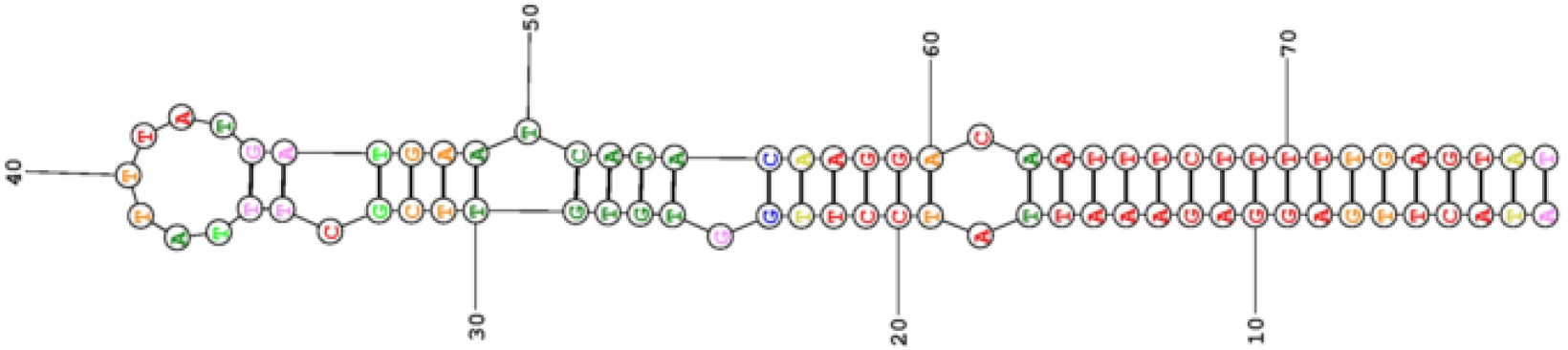
Homology model of the Homo sapiens miRNA-539 (MIR539). Source: Search result - http://rna.urmc.rochester.edu/RNAstructureWeb/

### Search for nucleotide sequence of miRNA-613

The main data source used for reconstructing the miRNA-613 was the nucleotide sequence file in FASTA format. The full-length nucleotide of miRNA-613 was obtained from the GenBank database under the identifier Homo sapiens miRNA-613 (MIR613), miRNA GenBank: NR_030344.1. The miRNA-613 was predicted to encode a 95 bp linear non-coding RNA, miRNA. All coding sequences were selected and exported as amino acids in FASTA format, using the annotation of the NCBI - Graphics. Homo sapiens miRNA-613 (MIR613), miRNA analysis is shown in **Figure 21**.

**Figure 21.**
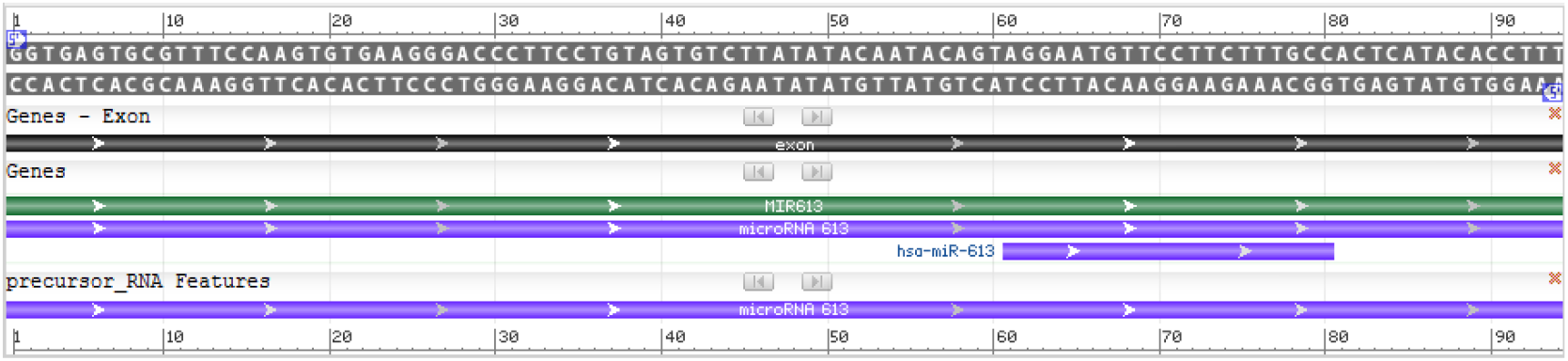
Homo sapiens miRNA-613 (MIR613) - *model-template aligment*. Source: http://www.ncbi.nlm.nih.gov/genbank/

### Building a molecular model of miRNA-613

Template search with FASTA format was performed against the RNAstructure template library, which were adjusted and optimized for alignment between Homo sapiens miRNA-613 (MIR613), miRNA, nucleotide and structural templates. On the basis of a sequence alignment between the miRNA-613 nucleotide and the template structure, a structural model for the target nucleotide was generated. Appraisal tools were utilized to calculate the trustworthiness of the obtained model. According to the described criteria, a model for the Homo sapiens miRNA-613 (MIR613), miRNA was generated (**Figure 22**).

**Figure 22.**
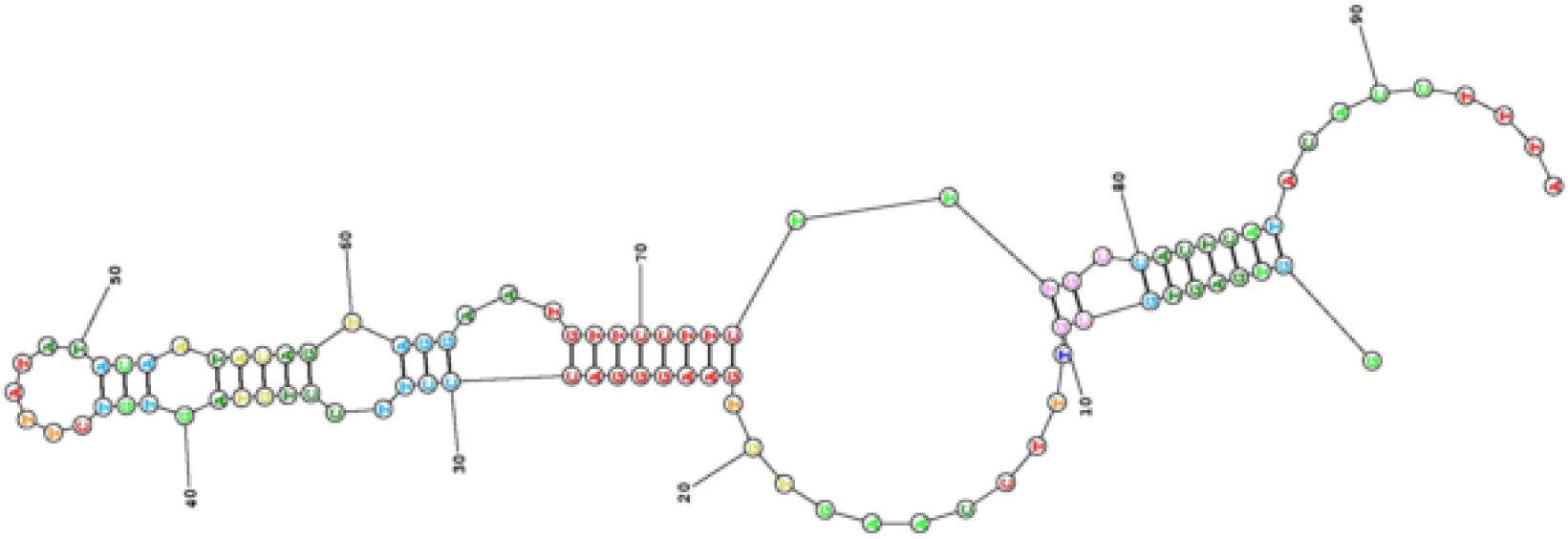
Homology model of the Homo sapiens miRNA-613 (MIR613). Source: Search result - http://rna.urmc.rochester.edu/RNAstructureWeb/

### Search for nucleotide sequence of miRNA-618

The reconstruction of the miRNA-618 was made from a nucleotide sequence file in FASTA format. The full-length nucleotide of miRNA-618 was obtained from the GenBank database under the identifier NCBI Reference Sequence: NR_030349.1. The Homo sapiens miRNA-618 (MIR618), miRNA was predicted to encode a 98 bp linear non-coding RNA, miRNA. All coding sequences were selected and exported as amino acids in FASTA format, using the annotation of the NCBI - Graphics. Homo sapiens miRNA-618 (MIR618), miRNA analysis is shown in **Figure 23**.

**Figure 23.**
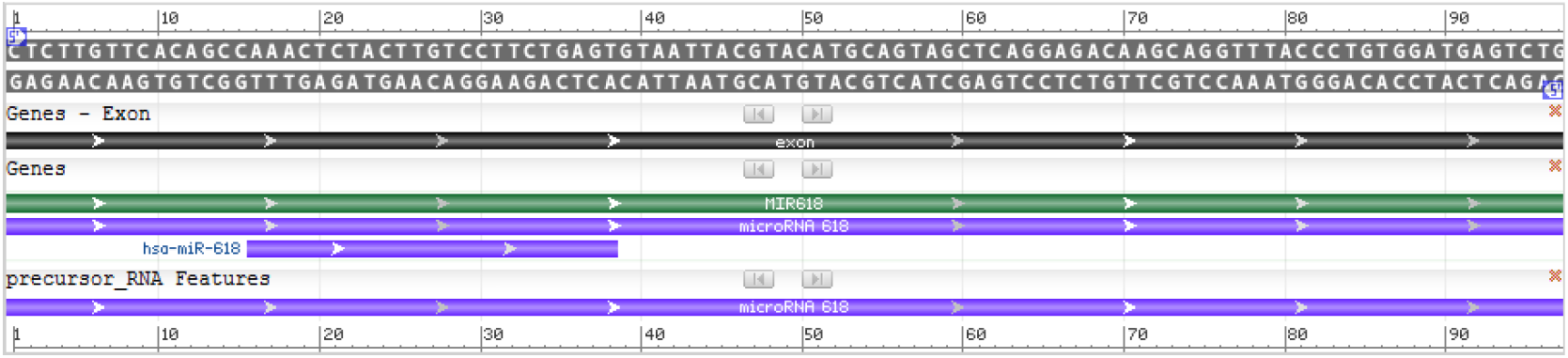
Homo sapiens miRNA-618 (MIR618) - *model-template aligment*. Source: http://www.ncbi.nlm.nih.gov/genbank/

### Building a molecular model of miRNA-618

Nucleotide sequences of miRNA-618 were obtain using FASTA format; modeling was conducted using the RNAstructure programs, which were adjusted and optimized for alignment between Homo sapiens miRNA-618 (MIR618), miRNA nucleotide and structural templates. On the basis of a sequence alignment between the Homo sapiens miRNA-618 (MIR618), miRNA nucleotide and the template structure, a structural model for the target nucleotide was generated. Appraisal tools were utilized to calculate the trustworthiness of the obtained model. Thus, using the RNAstructure programs automated comparative nucleotide modeling server, we constructed a homology model of the Homo sapiens miRNA-618 (MIR618), miRNA **(Figure 24)**.

**Figure 24.**
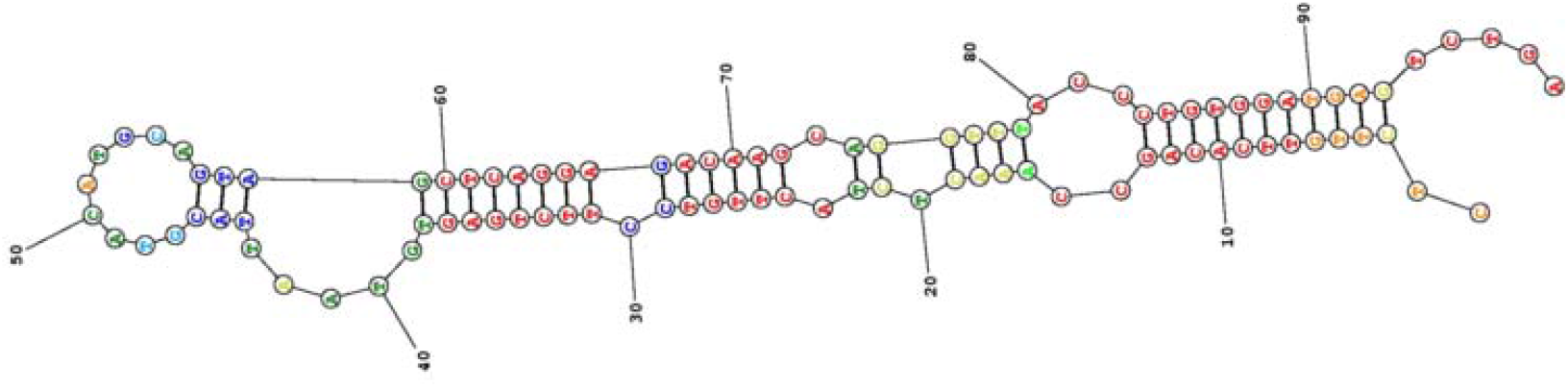
Homology model of the Homo sapiens miRNA-618 (MIR618). Source: Search result - http://rna.urmc.rochester.edu/RNAstructureWeb/

## DISCUSSION

The miRNAs are small non-protein-coding RNA that performs regulatory roles in several physiological and pathophysiological functions. In just over ten years, our knowledge of the structure and role of miRNA has significantly increased. In this study, we developed a tutorial on molecular modeling of 12 miRNAs already defined as miRNA inhibitors against miRNA over-expressed in thyroid cancer, realizing *in silico* projection of their molecular structures.

Studies have shown that miRNAs mutations or mis-expression are correlated with cancer and can act as tumor suppressors and oncogenes, being important in the diagnosis and treatment of thyroid cancer.^(5)^

The miRNA-101 is a small non-coding RNA that regulates several genes expression, is localized on chromosome 1 (1p31.3 position) and appears to be specific to the vertebrates, being validated in both human and mouse.^(6)^ MiRNA-101 has a stem-loop structure of about 70 nucleotides in length that is processed by the Dicer enzyme to form the 21-24 nucleotide mature miRNA, working as a tumor suppressor down-regulating a variety of malignancies including thyroid cancer and it low expression in papillary thyroid carcinoma (PTC) tissue was associated with lymph node metastasis and worse prognosis.^(7,8)^ Studies investigated the role of miRNA-101 in PTC and showed that miRNA-101 was considerably down-regulated in PTC specimen when compared with adjacent normal tissues, suggesting that miRNA-101 inhibited PTC growth via the down-regulation of Ras-related C3 botulin toxin substrate 1 (Rac1) expression.^(9)^ Following the same line of research, another study also demonstrated that miR-101 is significantly down-regulated in PTC tissue and thyroid cancer cell lines and that the down-regulated miR-101 is related with lymph node metastasis, suggesting that miRNA-101 operate as novel suppressor whose target is Rac1 gene, besides blocking thyroid cancer cell proliferation.^(10)^ Thus, the studies show new comprehension into the mechanism of inhibition of thyroid cancer by miR-101, suggesting the potential of miRNA-101 as a therapeutic target in thyroid cancer patients, in special of PTC.

Technological advances in post-genomic era have contributed to an expanding filling the databases and microarrays and other technologies have created a wealth of data for biologists, and the challenge facing scientists is to analyze and even to access these data to extract useful information. Several bioinformatics methods for miRNAs prediction have been developed, providing valuable understanding in the mechanisms of miRNAs transcriptional regulation. We researched the miRNA-101 sequences in the NCBI database using GenBank database under the identifier NCBI reference sequence, identifying all the nucleotide encoded in miRNA-101 and predicted their structure using domain analysis tools.

Recently, bioinformatics tools for the prediction of miRNAs have gained popularity because experimental studies for define miRNAs are unusual in their application. Nowadays, *in silico* evaluation of miRNA is based mostly on primary and secondary structure analysis. Reviewing the literature, we observed that none study with two-dimensional (2-D) structural model of the miRNA-101 was built. In our analysis, we carried out extensive the Nucleotide database that is a collection of sequences from several sources, including GenBank, RefSeq, TPA and PDB, and a 2-D model of miRNA-101 was built with the RNAstructure programs online.

The miRNA-126 is a short non coding RNA molecule, located within the 7th intron of epidermal growth factor-like protein 7 (EGFL7) gene of human chromosome 9 (9q34.3 position).^(11)^ miRNA-126 acts in angiogenesis control because is expressed only in endothelial cells by means of capillaries and larger blood vessels, and has miRNA-126 has previously been reported to be down-regulated in diverse types of cancer, including thyroid cancer.^(12,13)^ Recent study investigated the role and potential mechanism of miRNA-126 in tumorigenesis of PTC in vivo and in vitro, and demonstrated that its expression level was significantly down-regulated in PTC tissue and PTC cell lines, inhibiting cell proliferation, colony formations, migration and invasion, besides of promote cell apoptosis and cell cycle stop in G1 phase in vitro, and repress the tumor growth in vivo.^(14)^ other study investigated the role of miRNA-126 over vascular endothelial growth factor (VEGF)-A and their impact on angiogenesis in the several types of thyroid cancers, and the results demonstrated an under-expression of miRNA-126 compared to normal cells while VEGF-A were over-expressed, being promising the development of effective thyroid tumor-specific anti-angiogenic therapy.^(15)^ Reviewing the literature, we observed that in only one study a 2-D structural model of the miRNA-126 was built.^(16)^ In the study, the sequencing analysis of the miRNA-126 was evaluated and positioned in the 2-D structural model. In our study, the nucleotide analysis of miRNA-126 was performed in FASTA format, and the 2-D modeling used the RNAstructure program, which was adjusted and optimized for alignment between miRNA-126 and structural templates. Thus, in order to build a 2-D structural model of miRNA-126 we used the Nucleotide database sequence and a structural homology analysis strategy. The increased use of molecular models should promote advances in medicament engineering and could also facilitate the development of new therapies, as cited in previous reference.^(15)^

The miRNA-126-3p is an intronic miRNA, located in 7th intron of the EGFL7 gene in chromosome 9 (9q34.3 position), similar to miRNA-126, having as functions regulation of vasculogenesis and angiogenesis, besides playing an important role as either anti-oncogene in different cancer, because it behaves like tumor suppressor via the down-regulation of intron 7 of EGFL7.^(17)^ The role of miRNA-126-3p in thyroid cancer has been studied and suggests that his loss may be associated with thyroid cancer progression.^(18)^ Study to evaluate function of miRNA-126-3p in thyroid cancer cells showed that its expression was significantly lower in thyroid cancer, demonstrating that miRNA-126-3p has a suppressive function in thyroid cancer, being associated with the aggressive profile disease phenotype.^(19)^ We conducted NCBI-Nucleotide database searches using the miRNA domain as a query, which was identified by sequencing, and a high-quality structural model of miRNA-126-3p was built with the RNAstructure program.

The miRNA-141 is located in chromosome 12 (12p13 position) and belongs to the miRNA-200 family.^(20)^ The miRNA-141 is overexpressed in several human malignant diseases, and plays a dual role in tumorigenesis and being able to modulate cellular motility and control “stemness”.^(21)^ The miR-141 may distinguish PTC cell lines from a control thyroid cell line. Study to evaluation of miRNA-141 expression in thyroid cancer showed that miRNA-141 expression levels presented down-regulated in human thyroid cancer sample when compared with contiguous normal tissues, moreover its expression was considerably associated with clinical internship of tumor and presence of adjacent lymph nodes metastases.^(22)^ Other study investigated into the expression of miRNA-141 in a series of archival thyroid malignancies, and concluded that Hashimoto’s thyroiditis epithelium can be differentiated of PTC epithelium and control epithelium based in expression of miRNA-141.^(23)^ We built the 2-D molecular model of miRNA-141 using the homology modeling method based on the miRNA-141 sequence and the high-homology structure as the template with RNAstructure web server based on the nucleotide sequence of miRNA-141. The utilized computational system can have expressive value for expertise in thyroid cancer pathogenesis and may provide useful structural insights to facilitate rational drug design with personalized therapies.

The miRNA-145 is located on chromosome 5 (5q32-33 position) it is p53-regulated and has been implicated as both tumor and metastasis suppressor in multiple tumor types.^(24)^ Study to investigate the expression and the function of miRNA-145 in thyroid cancer and its potential clinical application as a biomarker showed that overexpression of miR-145 significantly down-regulate the thyroid cancer when compared with normal thyroid tissue resulting in reduction of cell proliferation, decrease VEGF secretion and E-cadherin expression, besides inhibited the phosphoinositide 3-kinase/threonine protein kinase, and thus also may serve as an additional biomarker for thyroid cancer diagnosis.^(25)^ Other study evaluate the expression of miRNA-145 in PTC and its potential function, and results demonstrated that the overexpression of miRNA-145 was lower in PTC tissues compared with adjacent normal specimens, and PTC cell line proliferation were suppressed, indicating that miRNA-145 might serve as a tumor suppressor gene of thyroid cancer, besides of propitiate a better understanding about the beginning and progression of PTC.^(26)^ A study evaluated *in silico* the miRNA structure through a 2-D computer model constructed with the UCSC Genome Browser 28 species conservation track, NCBI36 / hg18 assembly.^(27)^ Similarly, we built a 2-D structural model of Homo sapiens miRNA-145 (MIR145), based on the high-resolution structure, and this structural model was elaborated using RNAstructure.

The miRNA-146b belong the miR-146 family, is localized on chromosome 10 (10q24.32 position) that is transcribed into a precursor and produces two mature miRNAs (miRNA-146b-5p and miRNA-146b-3p).^(28)^ In PTC miRNA-146b-5p is highly expressed and positively associate to with degree malignancy.^(29)^ It has been demonstrated that miRNA-146b is suppressed in PTC, and was investigated the connection between the PTC recurrence risk and the expression of miRNA-146b and p27 protein miRNA in these tumors, and the results showed that the expression levels of miRNA-146b were high in cases of PTC with a elevated risk of recurrence.^(30)^ Study analyzed secondary structure of miRNA-146 using two computational approaches, MicroRNA.org (http://www.microrna.org) and targetscan (http://www.targetscan.org).^(31)^ In the present study, miRNA-146b nucleotides sequences were taken from GenBank database - NCBI, and we built the 2-D molecular model of miRNA-146b using the homology modeling method based on the miRNA-146b sequence with RNAstructure based on the nucleotide sequence of identifier Homo sapiens miRNA-146b (MIR146B).

The miRNA-206 is a vertebrate specific miRNA, and gene for human miRNA-206 is situated on chromosome 6 (6p12.2 position) in a bicistronic group.^(32)^ Has been proposed in preliminaries studies the tumor-suppressive function of miRNA-206 as well as has been reported the reduction of miRNA-206 expression in several solid neoplasms, among which are included the anaplastic thyroid cancer.^(33)^ A study demonstrated that the miRNA-206 inhibit proliferation and metastasis of anaplastic thyroid cancer by damage in myocardin family coactivator-A.^(34)^ A study showed the alignment and secondary structure of miRNA-206 constructed by the Mfold program (http://mfold.rna.albany.edu/).^(35)^ Similarly, we built a 2-D structural model of Homo sapiens miRNA-206 (MIR206), based on the high-resolution structure, and this structural model was elaborated using RNAstructure.

The miRNA-3666 is an RNA Gene family of miRNA class, and gene for human miRNA-3666 with chromosomal location 7 (7q31.1 position), and this gene have 1 transcript^36^. The function of miRNA-3666 in pathogenicity of thyroid cancer has been little studied. With respect to miRNA-3666, recent study analyzed the relationship between the miRNa-3666 levels and prognosis of patients with thyroid cancer, besides evaluated the levels of miRNA-3666 neoplastic thyroid tissue (NTS), observing that reduced levels of miR-3666 occurred in the NTS when compared to the adjacent non-tumor tissue, and also were associated with worse prognosis in survival of patients, yet the overexpression of miRNA-3666 presented an expressive growth inhibition of thyroid cancer cells.^(37)^ Searching in medical literature database observed that none scientific study with 2-D structural model of the miRNA-3666 was built. Searching in medical literature database, we observe that none scientific study demonstrating the 2-D structure model of the miRNA-3666 was built. In this study, the nucleotide sequence of miRNA-3666 was retrieved from NCBI sequence database, and this sequence was converted to FASTA format. The 2-D structure of miRNA-3666 was built using RNAstructure program. Thus, in absence of 2-D structures for most of the sequenced nucleotide, homology modeling experimentally forms the basis for the resolution of structure.

The miRNA-497 is a member of the miR-15 family with chromosomal location 17 (17p13.1 position). The miRNA-497 performs significant function in the pathogenesis of many diseases acting in signaling pathways in some pathologies including thyroid cancer, and annul cells multiplication, reduce the rate of S phase cells.^(38)^ Thus, miRNA-497 has demonstrated exercising the activity as a tumor suppressor. Study to investigate the function and supposed mechanism of miRNA-497 in thyroid cancer demonstrated that the miRNA-497 is a thyroid cancer tumor suppressor that acts blocking the oncogene brain-derived neurotrophic factor.^(39)^ Study analysed the secondary structure of miRNA-497 using the ClustalW command line version of the multiplatform sequence alignment and Mfold program (http://mfold.bioinfo.rpi.edu) was used to formation of secondary structure.^(40)^ Similarly, we built a 2-D structural model of Homo sapiens miRNA-497 (MIR497), based on the high-resolution structure, using the homology modeling method based on the miRNA-497 sequence and the high-homology structure as the template with RNAstructure web server based on the nucleotide sequence of miRNA-497.

The miRNA-539 is an RNA Gene family of miRNA class, with chromosomal location 14 (14q32.31 position). Studies have showed that miRNA-539 inhibit the tumor growth and metastasis in vivo, as well as has been demonstrating in vitro the inhibition, proliferation, migration, and invasion of neoplastic cells.^(41)^ Among the potential role of miRNA-539 draws attention to its tumor inhibitory activity on thyroid cancer.^(42)^ Study *in silico* employing computational calculation, it was verified 3’UTR of CARMA1 messenger RNA contains one miRNA-539 binding site, which would confirm the role of miRNA-539 in inhibition of thyroid cancer cell metastasis.^(43)^ Study in vitro using luciferase reporter assays proved that miR-539 binding to the 3’-UTR region of CARMA1 inhibit the expression of CARMA1 in thyroid cancer cells.^(42)^ In the analysis of the literature, we observed that none study with secondary structure model of the miRNA-539 was produced. We created a 2-D model of miRNA-539 using the RNAstructure program.

The miRNA-613 is a gene for human miRNA-613, and the locus of MiRNA-613 Gene is situated on chromosome 12 (12p13.1 position). The miRNA-613 have as main targets in tumors the KRAS, CDK4 and Frizzled7 molecular markers, also acts pleiotropic modulator in cancer cell multiplication and metastatic dissemination.^(44)^ Study to investigate the biological function and mechanism of miRNA-613 in the regulation of PTC development demonstrated that in vitro the overexpression of mirRNA-613 suppressed PTC cellular growth, migration and invasion, besides inhibit tumor growth in vivo intermediated by inhibition of SphK2 expression.^(45)^

The miRNA-618 is a short noncoding RNA molecule being situated on chromosome 12 (12q21.31 position), control a series of cancer-related gene networks and pathways, and whose deregulation is associated to malignancies neoplasms including thyroid cancer.^(46)^ Study demonstrated that the miRNA-618 had a function of down-regulated on thyroid cancer cell lines,^(47)^ and other study evaluated the expression, role and the mechanism of miRNA-618 in human thyroid cancer cells, confirmed those findings, proving that functionally the forced expression of miRNA-618 represses the cellular multiplication, and from mechanical point of view the miRNA-618 overexpression induces the inhibition of PI3K/Akt signaling pathway in Human TPC-1 cell line.^(46)^ Schematic of secondary structure for miRNA-618 was presented in study at where the structure image was adapted from the Vienna RNA secondary structure prediction algorithm (http://www.tbi.univie.ac.at/RNA/).^(48)^ We carried out extensive the Nucleotide database that is a collection of sequences from several sources, including GenBank, RefSeq, TPA and PDB, and a 2-D model of miRNA-618 was built with the RNAstructure programs online.

## CONCLUSION

The structure and function of miRNAs are determined by their nucleotides sequences and high resolution structure prediction methods make possible to identify the location of binding sites on nucleotides of fundamental importance for applications clinical and pharmacological aspects, mainly in oncological therapy.

In this study we show *in silico* secondary structures projection of selected of 12 miRNA inhibitors against miRNA over-expressed in thyroid cancer through computational biology.

